## Supplementary information for "Inhibitor-based modulation of huntingtin aggregation mechanisms mitigates fibril-induced cellular stress"

**This PDF file includes:**

- Extended Methods
- Supplementary Figures 1-10
- Supplementary Tables 1-4
- References cited in the Supplementary Information

### Extended Methods

#### Protein expression and purification

The maltose-binding-protein (MBP)-fusion proteins MBP-Q32-HttEx1 and MBP-Q44-HttEx1, featuring HttEx1 with 32 and 44 consecutive glutamine residues within the polyQ segment, were sub-cloned into a pMALc5x plasmid (for Q32) or into pMALc2x plasmid (Q44) by Genscript (Piscataway, NJ) as previously described (1). The fusion protein was overexpressed in E. coli BL21(DE3) pLysS cells (Invitrogen, Grand Island, NY). The cells were grown in 2L LB medium with ampicillin and chloramphenicol at 37°C and 200 r.p.m. until an optical density (OD600) of 0.80. A temperature ramp from 37°C to 18°C over 30 mins at 200 r.p.m. was applied to prepare for induction. Proteins for study by ssNMR were uniformly $^{13}$C, $^{15}$N labeled with $^{13}$C-D-glucose and $^{15}$N ammonium chloride (Sigma-Aldrich) during overexpression, using a previously described protocol optimized for isotopic labeling (2). Protein expression was induced by adding 0.6mM IPTG (Sigma-Aldrich). The protein was over-expressed at 18°C for 16 hours, after which the cells were pelleted at 5250xg for 20mins. The cells were resuspended in Phosphate Buffered Saline (PBS) buffer (11.9mM phosphates, 137mM sodium chloride and 2.7mM potassium chloride), pH 7.4. The resuspended cells were kept on ice, followed by the addition of 1mM phenylmethanesulfonyl fluoride (Sigma-Aldrich), and 0.5mg/ml of lysozyme (Sigma-Aldrich). The cells were burst using sonication (VCX130 Vibra-cell sonicator, Sonics & Materials, Inc), applying 70% amplitude for 20 mins with alternation of 10s pulses and breaks of 10s. Cell debris was removed by centrifuging at 125,000xg for 45mins. The soluble protein was filtered using 0.45µm syringe filters and purified using a HisTrap HP nickel column (GE Healthcare) with an imidazole gradient. The purified protein was exchanged into an imidazole-free PBS buffer using dialysis membrane of 12400 MWCO (Sigma-Aldrich). The concentration of fusion protein was determined by its average absorbance (n=3) at 280nm, as measured in a 50µl quartz cuvette with a 10mm path length by a JASCO V-750 spectrophotometer. The extinction coefficient of the fusion protein is estimated to be 66,350 $M^{-1}cm^{-1}$ (from the protein sequence by the ProtParam tool by ExPASy (3)). The protein purity and molecular weight were verified by ESI-TOF MS and SDS-PAGE (12%) as described previously (1).

**Fibril formation**

The purified fused protein was cleaved by treating it with Factor Xa (FXa) (Promega, Madison, WI) at room temperature to release the HttEx1 (HttEx1:FXa = 400:1 molar ratio). The progression of cleavage and aggregation were monitored by SDS-PAGE (Bio-Rad mini protein Precast TGX Gels 12%) and TEM. The fusion protein was cleaved and allowed to aggregate over 4 days, after which the HttEx1 fibrils were pelleted down at 3,000xg for 20 min. The supernatant was discarded, and pelleted fibrils were washed at least three times with PBS buffer. For MAS NMR-analyzed samples 1-5, the fusion protein was used in a uniformly $^{13}$C,$^{15}$N-enriched form (see below and Supplementary table 1). For Sample 5, 71μM of MBP-Q32-HttEx1 protein was cleaved by treating it with FXa at room temperature to release the HttEx1 (HttEx1:FXa = 400:1 molar ratio). The fusion protein was allowed to aggregate over 4 days, and then the fibrils were pelleted down at 3000xg for 20 mins. The fibrils were washed three times with PBS buffer and then re-suspended in PBS. Then, these pre-formed fibrils were treated with 23.6μM of curcumin in PBS for a 1:0.33 protein to curcumin molar ratio. For the TEM studies, the sample was collected at specific time intervals (24, 48, 72 hours). For ssNMR, these fibrils with curcumin were packed in the rotor, and the sample was measured at specific time intervals (24, 48, 72, 96 hours).

**ThT assay**

Aggregates were re-suspended by aspiration. A 1mM Thioflavin T (ThT) stock was prepared in DMSO, which was further diluted in PBS buffer. The final concentration of ThT for the assay was 15μM. Curcumin stock was prepared (1mM) in DMSO and diluted in PBS buffer for spectroscopy at a final concentration of 8.5μM, 12μM, and 20μM, using a concentration of 61μM of MBP-Q32-HttEx1 protein. The FXa was added in 400:1 molar ratio (HttEx1:FXa). Samples were mixed gently with a pipette and thereafter measured in black polystyrene, clear bottom, 96-well plates (Corning, U.S.A) on TECAN Spark 10M microplate reader. All fluorescence measurements (3 independent replicates) were taken at 25ºC. Samples were excited at 442nm and emission was recorded between 482-492nm, using excitation and emission bandwidths of 10nm. The plate was shaken for 5s before recording the emission for each measurement. The emission was recorded every 15 minutes at 490nm and was used for plotting the fluorescence kinetics curves (Figure 2), based on the subtraction of the signal from the control (PBS buffer with ThT). Plotted data were normalized to the maximum observed signal. For studying the aggregation kinetics of Q44-HttEx1, the same protocol was followed with a HttEx1 protein concentration 67.5μM and curcumin concentrations of 10μM and 20μM. The aggregation curves in Figure 2a-b were plotted in Origin 8.1 software, and represent the average values of data measured in triplicates. The aggregation curves with error bars are shown in Supplementary figure 2a-c.

**Transmission electron microscopy**

Transmission electron microscopy (TEM) was performed on mature Q32-HttEx1 and Q44-HttEx1 protein fibrils. 61μM of the fused Q32-HttEx1 and 50μM of Q44-HHttEx1 protein was cleaved by treating it with FXa (Promega, Madison, WI) at room temperature to release the HttEx1 (HttEx1:FXa = 400:1 molar ratio). The protein fibrils which were allowed to aggregate at room temperature for 96 hours in the absence and presence of different concentrations of curcumin. For the studying the effect of curcumin on the 71μM of the Q32-HttEx1 pre-formed fibrils, these fibrils were treated with 23.6μM of curcumin (i.e. protein: curcumin; 1:0.033 molar ratio). For the imaging studies, the fibrils were suspended in MiliQ water. The fibril samples were deposited on plain carbon support film on 200 mesh copper grids (SKU FCF200-Cu-50, Electron Microscopy Sciences, Hatfield, PA) after glow-discharge for 0.5 -1 min. The excess MiliQ was removed by blotting. The negative staining agent used was 2% (w/v) uranyl acetate. The stain was applied immediately after blotting for 0.5 - 1 min. The excess stain was removed by blotting and the grid air dried. The images were recorded on a Philips CM120 electron microscope operating at 120kV. Images were recorded on a slow scan CCD camera (Gatan). The fibril widths were measured transverse to each fiber axis using the straight free-hand tool of Fiji (4). Each measurement spanned the length of the negative stained area of the fibril with similar contrast The width measurements are assumed to reflect the polyQ core, as the stain accumulates on the flanking domains (1). In images with low resolution, the fibril diameter was determined in regions with the clearest defined boundaries. Three measurements were obtained per fibril, except for fibrils where the width varied significantly. In selected cases, fibril widths were verified on isolated and vertically aligned fibrils using the Plot Profile tool of Fiji, which plots the average grayscale intensity values across the fibril axis.

**Cell culture**

Mouse HT-22 hippocampal cell lines and Lund human mesencephalic cells (LUHMES)-differentiated neuronal cells were kindly provided by Prof. Culmsee, University of Marburg, Germany. HT22 cells were cultured in Dulbecco's Modified Eagle Medium (Gibco, ThermoFisher Scientific, Landsmeer, The Netherlands) supplemented with 1% pyruvate (ThermoFisher Scientific), 10% fetal bovine serum (GE Healthcare Life Sciences, Eindhoven, the Netherlands), and 100U/mL penicillin-streptomycin. LUHMES cells were cultured in Advanced DMEM/F12 (Gibco), supplemented with 200 mM L-Glutamine (Gibco), 1x N2 Supplement (Gibco), 100 µg/ml FGF (PeproTech, Cranbury, USA), and 1x penicillin/streptomycin solution (Gibco). LUHMES cells were differentiated to dopaminergic neurons before the treatments as described before (5, 6). Cells were maintained at 37°C and 5% $CO_2$. Cells were used for experiments for at most 10 passages after thawing and regularly checked for Mycoplasma infection.

**Cell toxicity assay:** Cell toxicity was assessed using the CytoTox-Glo™ Cytotoxicity Assay (Promega). HT22 cells were seeded in white-walled 96-well plates at a density of 8 × $10^3$ cells per well in 100 µL of medium and incubated at 37°C with 5% $CO_2$ for 24 hours. The cells were then treated with 5, 15, or 25 µM of pre-formed Q32-HttEx1 fibrils in medium for 72 hours, with untreated wells serving as negative controls. After treatment, 50 µL of CytoTox-Glo™ Reagent was added to each well. The plates were briefly mixed, incubated for 15 minutes at room temperature, and luminescence was measured to quantify cell death (read 1). Then, 50 µL of Lysis Reagent was added to lyse all cells, followed by another 15-minute incubation. Luminescence was measured again to determine total cell activity (read 2). Measurements were taken using a Synergy H1 Multi-Mode reader (Biotek, Winooski, USA). Live cell activity was determined according to the equation: Live Cell Activity = Total Luminescence (read2) − Experimental Dead Cell Luminescence (read1). The data was normalized by the average of the cell activity of the control group.

**Small angle X-ray measurements (SAXS)**

63μM of MBP-Q32-HttEx1 was cleaved using Factor Xa protease in molar ratio (HttEx1:FXA = 400:1) and aggregated in presence and absence of curcumin (1:0.33) at room temperature. After 96 hours, the fibrils were collected as described in methods earlier. The fibrils were centrifuged at 2400xg, and the supernatant (PBS) was removed such that to make the final concentration of the fibrils to 2mM for the SAXS experiment. The experiments were performed at the multipurpose X-ray instrument for nanostructured analysis (MINA) at the University of Groningen. The instrument is equipped with a high flux rotating anode source using radiation from a Cu anode and delivering X-ray photons with a wavelength $\lambda$ = 0.15413 nm (E = 8 keV). The SAXS patterns have been acquired using a solid-state noiseless Pilatus 300k detector (Dectris) placed 3.1m away from the sample. The coordinates of the transmitted X-ray beam and the angular range probed by the measurements were calibrated using a standard diver behgenate powder (NIST). The 2D SAXS patterns were reduced to the 1D SAXS profiles using the Fit2D and Origin software, after proper correction for the difference in sample absorption and subtraction of the background (PBS buffer). Samples were measured in 1.5mm glass capillaries and at ambient temperature (23°C). The SAXS curves are plotted as I(q) vs q in the log-log scale, where $q = (4\pi sin\theta)/\lambda$ is the modulus of the scattering vector and $2\theta$ is the scattering angle.

**SAXS data analysis**

The SAXS curves have been analysed using the model for a concentrated ensemble of long rod-like objects characterised by length L and radius r. The general equation for the fitted model in the framework of the monodisperse approximation is:

$$I(q) = A\left[\int_0^{\infty} P(q,r,L)N(r)dr\right]S(q,d,\nu) + I_{bkg}$$

where $P(q,r,L)$ is the function called form factor, describing the scattering for the rod-like objects, and $S(q,d,\nu)$ is the function called the structure factor, describing the lateral packing of the rod-like objects, with average interfibrillar distance d and interaction parameter $\nu$. The fibrillar cross-sectional dimension is subjected to a log-normal size distribution $N(r)$, characterised by an average dimension $\bar{r}$ and width $\sigma$. The prefactor A is a constant which includes the number density of objects (i.e. concentration) and the squared of the average electron contrast between the Q32-HttEx1 fibrils and the surrounding media. The background intensity $I_{bkg} = B/q^n + C$ takes into account all atoms and molecules which are not contained in the measured background and are not included in the assembled fibrillar domains.

As form factor we used the standard equation for long cylindrical objects (7). Since the length L of the rod-like objects is outside the probed range, its value was kept fix to 1000 nm during fitting. For the structure factor we used the expression derived for the Polymer Reference Interaction Site Model (PRISM) (8, 9). In this model, the interaction parameter $\nu$ is associated with the second virial coefficient describing the pair interaction between adjacent cylinders. All equations are implemented in the SASfit software (10) which was used to fit the SAXS 1D profiles. The fitted parameters are thus A, r, d, $\nu$, B, C and n (Supplementary Table 3).

In addition to the model above, the data have been fitted also using a model previously reported by Perevozchikova et al. for single huntingtin fibrils (11). The average fibrillar dimensions obtained using this model and characterized by a radius of cross-sectional radius of $R_{g1}$ are in close agreement with the ones obtained by our model. The fitted curve using the model by Perevozchikova et al. for the Q32 sample prepared without curcumin is reported in Fig. 2o in the main manuscript as dashed line, while the values for the fitted parameters are reported in supplementary table 4 for fibrils prepared without and with curcumin.

**MAS solid-state nuclear magnetic resonance**

All solid-state NMR experiments were acquired on a Bruker Avance NEO 600 MHz spectrometer, equipped with a 3.2 mm MAS probe with an HCN Efree coil (Bruker Biospin). Isotopically labeled Q32-HttEx1 fibrils in the absence and presence of curcumin were prepared and then packed by pelleting a hydrated suspension of purified protein fibrils into 3.2mm zirconia thin wall MAS rotors (Bruker Biospin, Billerica, MA). This sedimentation process was done using a home-built ultracentrifugal packing device under centrifugation at ~130,000xg in a Beckman Coulter Optima LE-80K ultracentrifuge equipped with an SW-32 Ti rotor (12). A summary of sample details is provided in Supplementary Table 1 below, referring to the individual preparations as Samples 1 through 5. Samples 1 and 2 were prepared together (referred to as Batch 1), being Q32-HttEx1 (control; Sample 1) and Q32-HttEx1 with curcumin (1:0.33). Samples 3 and 4 represent separate Batch 2, of otherwise identically treated Q32-HttEx1 (control) and Q32-HttEx1 with curcumin (1:0.33). In case of sample 5, the labeled Q32-HttEx1 fibrils were treated with curcumin (1:0.33) after preparation of the fibrils. Caps were sealed to the rotor with epoxy glue to ensure stable sample hydration. Samples were studied by MAS ssNMR in a hydrated and unfrozen state. All the 1D experiments for the Batch 1 and Batch 2 fibril samples were recorded with a spinning frequency of 10kHz, 1024 scans, and at 275K. The 2D $^{13}C$-$^{13}C$ DARR (13) experiment was performed with a 25ms mixing time with 40 scans at 13kHz spinning rate. The 2D $^{13}C$-$^{13}C$ INEPT- TOBSY ($P9^1_3$) experiment was performed at 8.33kHz MAS with 32 scans (14). More experimental details are in Supplementary Table 2. SSNMR peak integration analysis was done on the $^{13}C$ cross-polarization spectra obtained for batch 1 and batch 2, Q32-HttEx1 (control) and Q32-HttEx1 with curcumin (1:0.33) fibrils. Spectra were processed with NMRPipe (15) and peaks were integrated between the mentioned chemical shifts ranges with a python script (available from nmrglue website (16)). For batch 1 fibrils, the peak area for PC$\alpha$ was selected from 68.31 ppm to 60.68 ppm; for QC$\alpha$ - 60.68 ppm to 53.40 ppm and for PC$\delta$ - 53.40 ppm to 47.78 ppm. For batch 2 fibrils, the peak area for PC$\alpha$ was selected from 66.70 ppm to 58.99 ppm; for QC$\alpha$ - 58.99 ppm to 51.79 ppm and for PC$\delta$ - 51.79 ppm to 46.01 ppm. The obtained values were normalized by dividing them by the maximum intensity corresponding to the QC$\alpha$ peak and visualized in Microsoft Excel. Error bars represent an estimate of the noise.

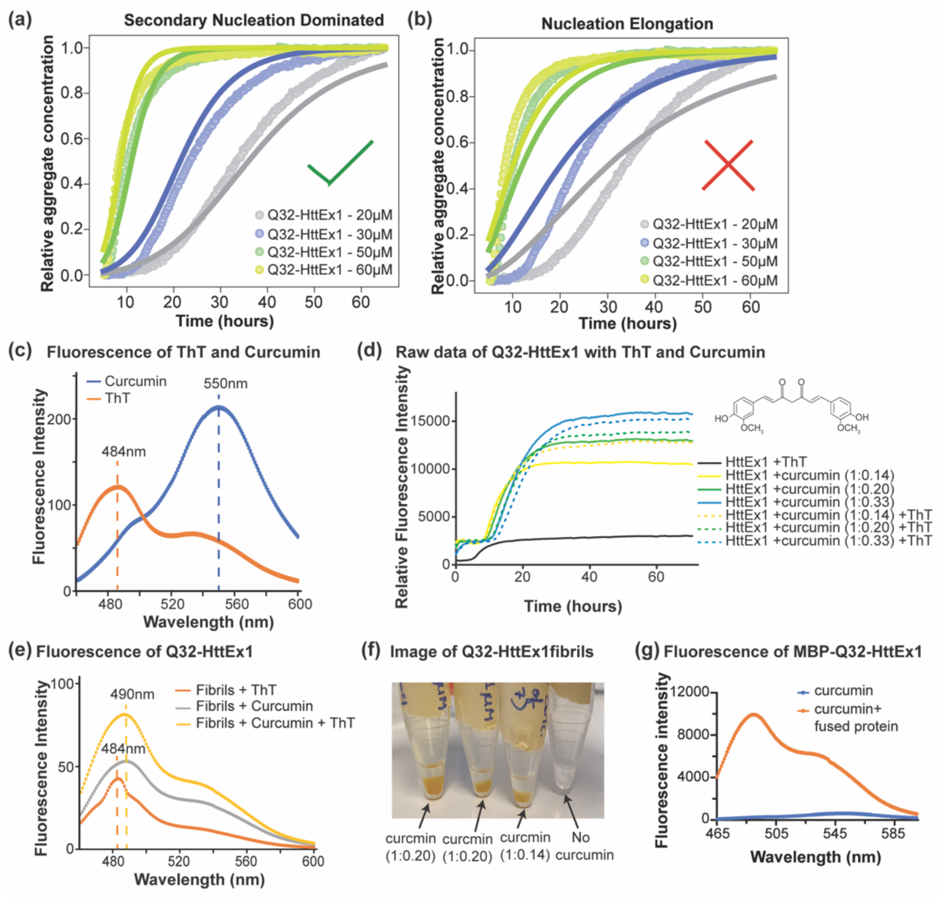


**Supplementary Fig. 1. Fluorescence data of ThT and curcumin.** (a,b) ThT kinetics data for Q32-HttEx1 aggregation at different concentrations (20μM, 30μM, 50μM and 60μM) were analyzed and plotted in Amylofit (17). Two kinetic models are shown: (a) secondary nucleation dominated, and (b) nucleation elongation (without secondary nucleation). The fitting suggests that the aggregation proceeds predominantly via secondary nucleation. The former mechanism better fits the data. The ThT data were measured in technical triplicates and the mean of the triplicates was plotted. (c) Fluorescence of curcumin and ThT in buffer when excited at 442nm. The dotted lines show the emission maxima for ThT at 484nm and for curcumin at 550nm. (d) Q32-HttEx1 (61μM) aggregation was monitored by ThT fluorescence in the presence and absence of curcumin at various sub-stoichiometric molar ratios, as indicated. Solid lines include samples without ThT, but with curcumin. Dashed lines show the same protein: curcumin ratios with also 15μM ThT present. Note that the fluorescence signal in presence of curcumin is not only showing a delayed increase, but also reaches a much higher fluorescence intensity. The increased intensity stems from the binding of curcumin to the formed HttEx1 fibrils (see panel f), which results in the immobilization of the normally flexible molecule (inset top right) and causes a dramatic increase in curcumin fluorescence. Inclusion of ThT in the sample yields a fluorescent signal intensity that is a combination of the two fluorophores but dominated by the curcumin signal. (e) Fluorescence of Q32-HttEx1 fibrils (30μM) with ThT (15μM) and curcumin (10μM) excited at 442nm. The dotted lines show the emission maxima of ThT bound to fibrils at 484nm and curcumin and ThT bound to the fibrils at 490nm. (f) Photograph of HttEx1 fibrils formed in presence and absence of curcumin, after centrifugation. The fibrils in the pellet display a clear coloring due to curcumin bound to the fibrils. (g) Fluorescence of 10μM curcumin alone compared to that of 10μM curcumin in presence of MBP-Q32-HttEx1 fusion protein (50μM), in PBS and excited at 442nm. Note that the fluorescence intensity units of different panels should not be compared, due to different settings of the spectrofluorometer.

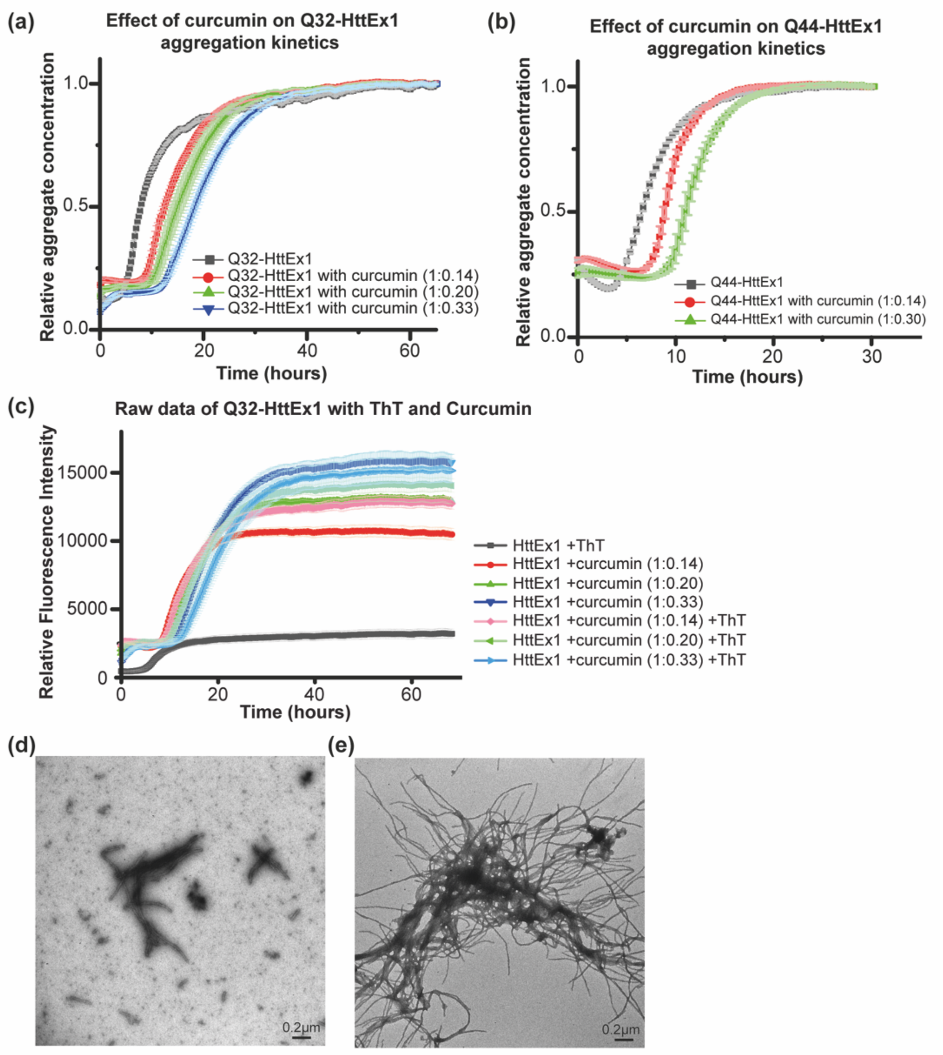


**Supplementary Fig. 2. Fluorescence data with error bars and EM analysis showing more bundled fibrils formed in presence of curcumin.** (a) Q32-HttEx1 (61μM) aggregation curves (also in figure 2a) measuring fluorescence in the presence and absence of curcumin (molar ratio indicated). (b) Q44-HttEx1 (67.5μM) aggregation fluorescence curves (also in figure 2b) in the presence and absence of curcumin. The curves indicate the mean of triplicate values with error bars (standard deviation). (c) Q32-HttEx1 (61μM) fluorescence aggregation curves (also in Supplementary Figure 1d) in the presence and absence of curcumin at various sub-stoichiometric molar ratios, as indicated. Dark red, green and blue lines include samples without ThT, but with curcumin. Light red, green and blue lines show the same protein:curcumin ratios with also 15μM ThT present. Note that the fluorescence signal in presence of curcumin is not only showing a delayed increase, but also reaches a much higher fluorescence intensity. Inclusion of ThT in the sample yields a fluorescent signal intensity that is a combination of the two fluorophores, but dominated by the curcumin signal. (d) Negative stain TEM micrograph of Q32-HttEx1 fibrils prepared *in-vitro* at room temperature. (e) TEM micrograph of Q32-HttEx1 fibrils formed in presence of curcumin (1:0.33), prepared *in vitro* at room temperature. Note the more bundled fibrils in comparison to the fibrils formed in absence of curcumin. These observations are highly reproducible and have been repeated multiple times.

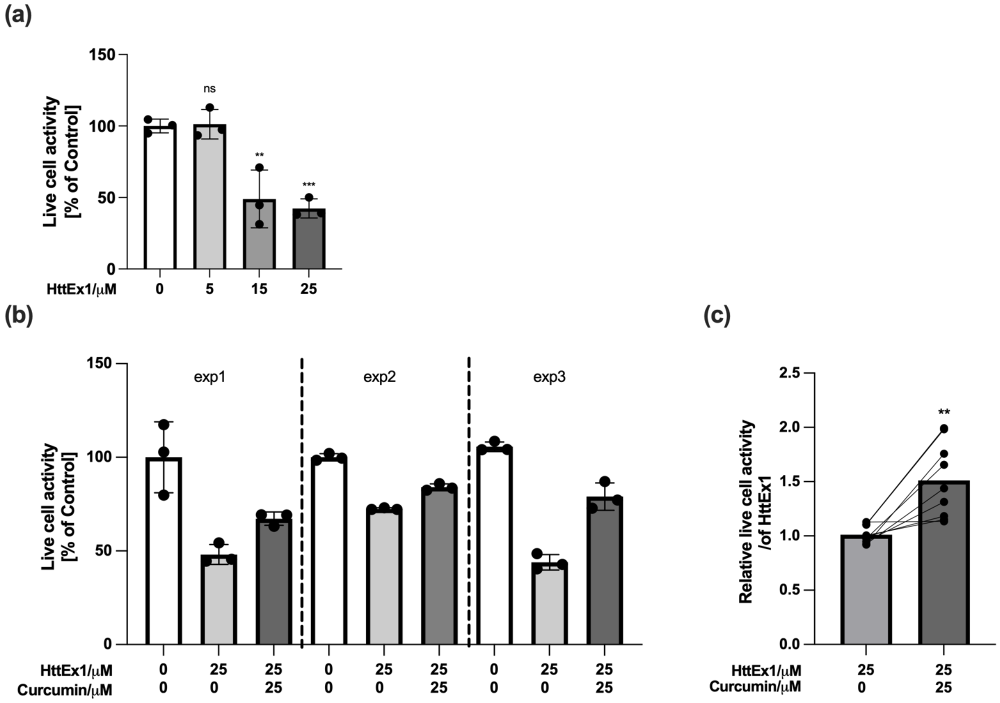


**Supplementary Fig. 3. Assays of cytotoxicity caused by HttEx1 fibrils in HT22 cells. (**a) Effect of Q32-HttEx1 fibrils on HT22 cells. Live cell activity in HT22 cells was measured using the CytoTox-Glo™ Cytotoxicity Assay, which measures the release of (cytosolic) protease activity from compromised cells, as a measure of cell death. HT22 cells were treated with varying concentrations of Q32-HttEx1 fibrils (5, 15, or 25μM) for 72 hours. Live cell activity was quantified by subtracting the dead cell activity from the total cell activity, measured after cell lysis (see Extended Methods above). These data are obtained from 3 independent experiments with 9 technical replicates each. The error bars indicate mean with SD. (b) Effect of curcumin-inhibited Q32-HttEx1 fibrils on treated cells. Live cell activity from three independent experiments is shown for HT22 cells treated with 25μM Q32-HttEx1 fibrils or with 25μM Q32-HttEx1 fibrils obtained after curcumin inhibition, for 72 hours. Control samples are cells not exposed to fibrils. Error bars indicate mean with SD (c) Relative rescue effect of curcumin inhibition. Relative live cell activity was calculated from data in panel (b), with each dot representing a single technical replicate (9 technical replicates per condition across three biological replicates). Data were normalized to the mean of the 25μM Q32-HttEx1 fibril condition to illustrate the relative rescue effect associated with curcumin inhibition. Statistical significance was determined using an unpaired t-test ($p < 0.01$).

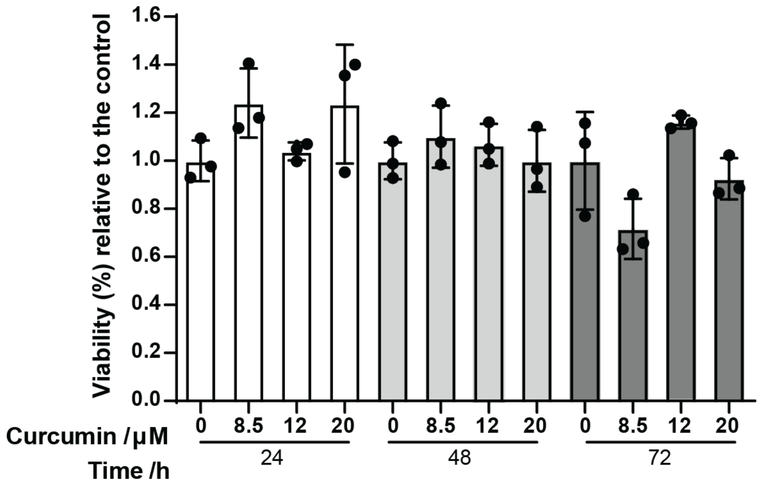


**Supplementary Fig. 4**. **Cell viability of mouse hippocampal HT22 cells upon exposure to varying concentrations of curcumin (in absence of any HttEx1 fibrils).** HT22 cells were seeded in p96 wells at a density of $9.10^3$ cells/well. After 24 hours, the cells were treated with 0, 8.5, 12 or 20 µM curcumin diluted in culture medium and incubated for 24, 48 or 72 hours at 37$^o$C and 5% $CO_2$. Cell viability was assessed through the MTT reduction assay. Absorption values are normalized to the average of the untreated control.

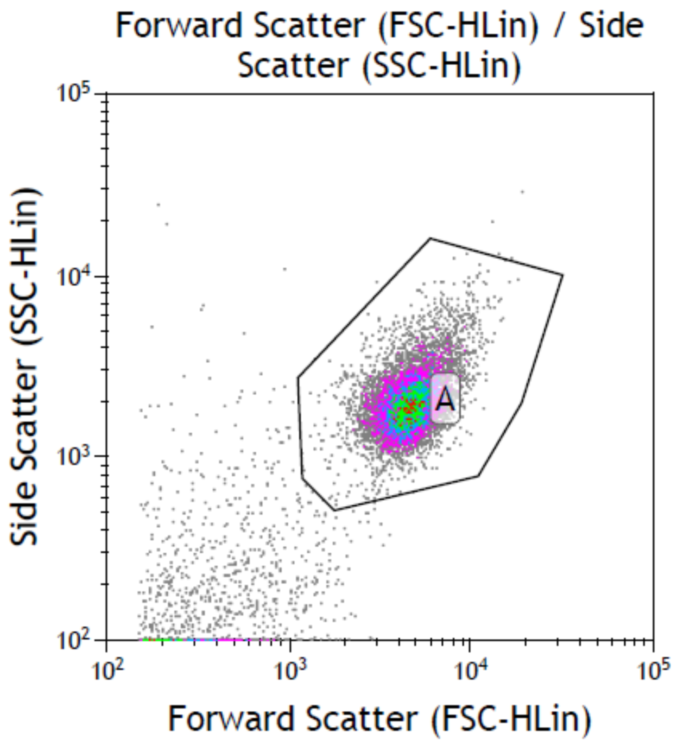


**Supplementary Fig. 5.** Cytometry gating strategy. A representative image of the population of interest (A). The selection of the interest population was based on particle size in the SSC versus FSC plot, aiming to exclude debris. The same gate was applied to all conditions and the mean intensity of fluorescence in the FITC channel was evaluated. The population of interest was greater than 75% in all analyzed conditions.

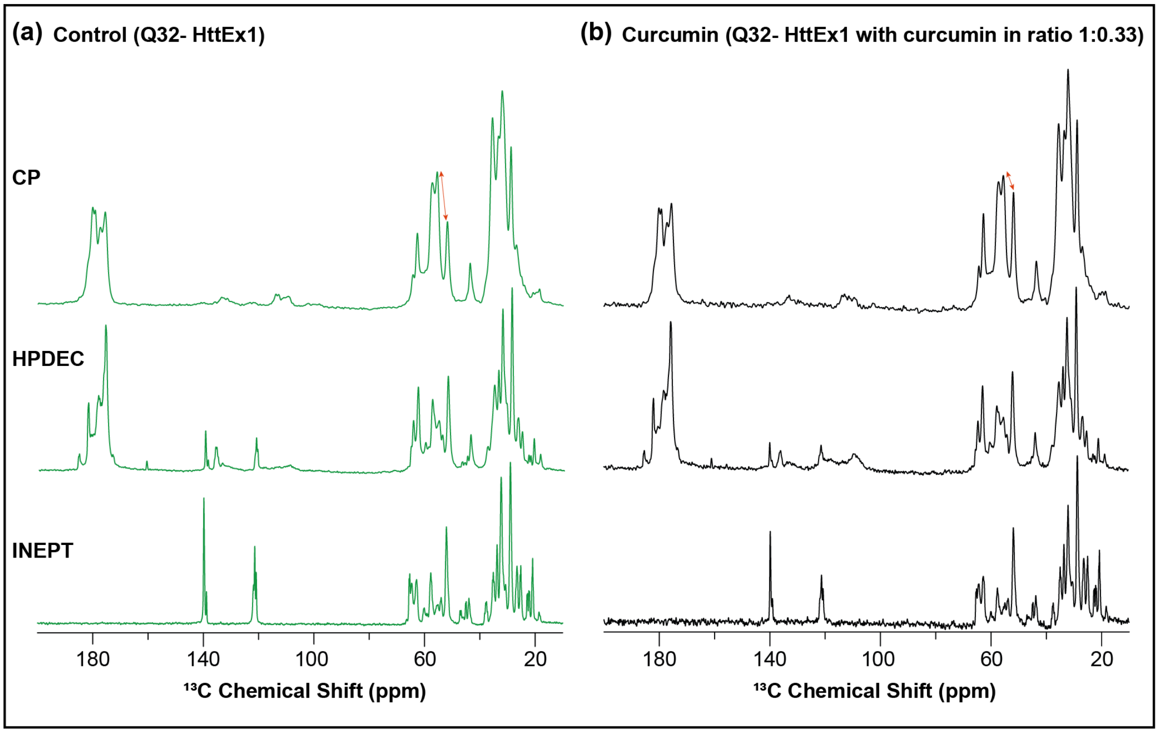


**Supplementary Fig. 6. MAS ssNMR comparison of Q32-HttEx1 and Q32-HttEx1 with curcumin (1:0.33) (Batch 1 fibrils).** (a) 1D CP $^{13}C$ spectrum (top), 1D direct excitation (HPDEC) $^{13}C$ spectrum (middle) and 1D INEPT $^{13}C$ spectrum (bottom) of Q32-HttEx1 fibrils. (b) 1D CP $^{13}C$ spectrum (top), 1D HPDEC $^{13}C$ spectrum (middle) and 1D INEPT $^{13}C$ spectrum (bottom) of Q32-HttEx1 with curcumin (1:0.33) fibrils. The CP spectra feature signals from rigid and partly immobilized parts of the structure, while the INEPT data show only highly flexible residues. Note the presence of strong signals from the C-terminal His tags in the region of 120-140 ppm (aromatic region) of the INEPT spectra, which shows the highly flexible C-termini being exposed on the fibril surface. The orange arrows in the CP spectra shows the signal intensity between the glutamine and proline signals. Thus, showing the increase in the proline signals for the fibrils formed in presence of curcumin.

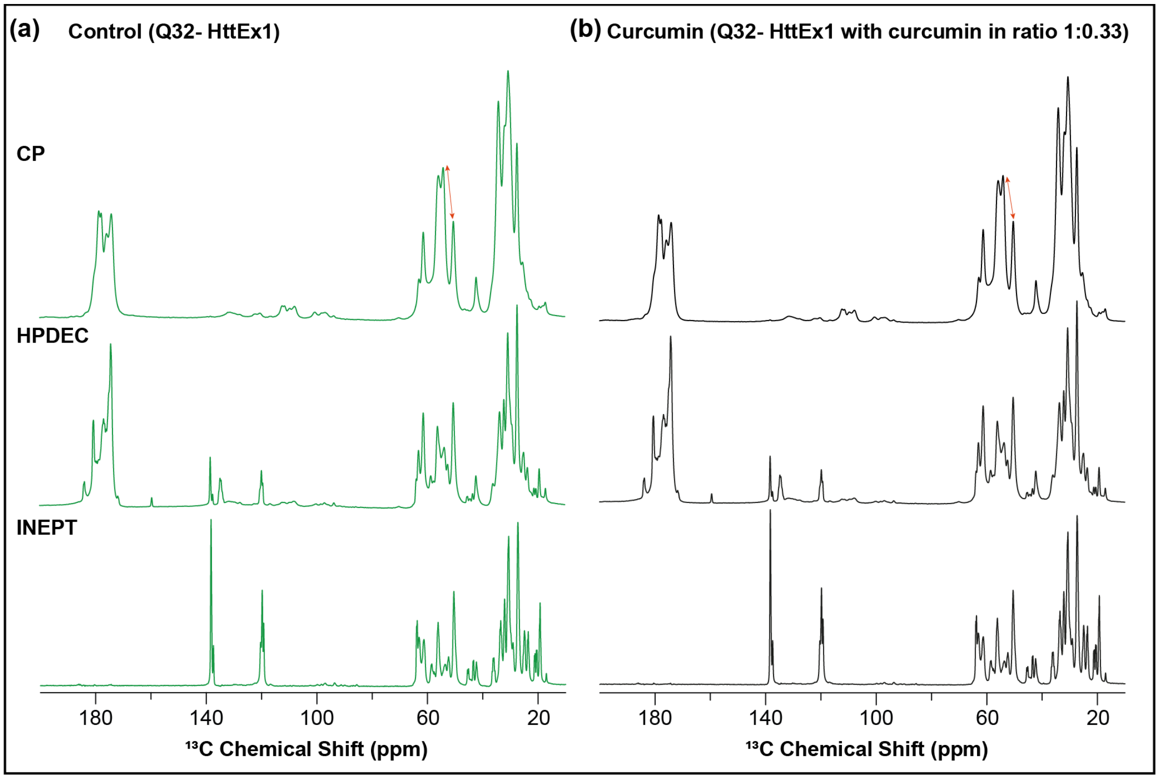


**Supplementary Fig. 7. MAS ssNMR comparison of Q32-HttEx1 and Q32-HttEx1 with curcumin (1:0.33) (Batch 2).** (a) 1D CP $^{13}C$ spectrum (top), 1D HPDEC $^{13}C$ spectrum (middle) and 1D INEPT $^{13}C$ spectrum (bottom) of Q32-HttEx1 fibrils. (b) 1D CP $^{13}C$ spectrum (top), 1D HPDEC $^{13}C$ spectrum (middle) and 1D INEPT $^{13}C$ spectrum (bottom) of Q32-HttEx1 with curcumin (1:0.33) fibrils. The CP spectra feature signals from rigid and partly immobilized parts of the structure, while the INEPT data show only highly flexible residues. Note the presence of strong signals from the C-terminal His tags in the region of 120-140 ppm (aromatic region) of the INEPT spectra, which shows the highly flexible C-termini being exposed on the fibril surface. The orange arrows in the CP spectra shows the signal intensity differences between the glutamine and proline signals (see also Supplementary Fig. 8 below). Thus, showing the increase in the proline signals for the fibrils formed in presence of curcumin.

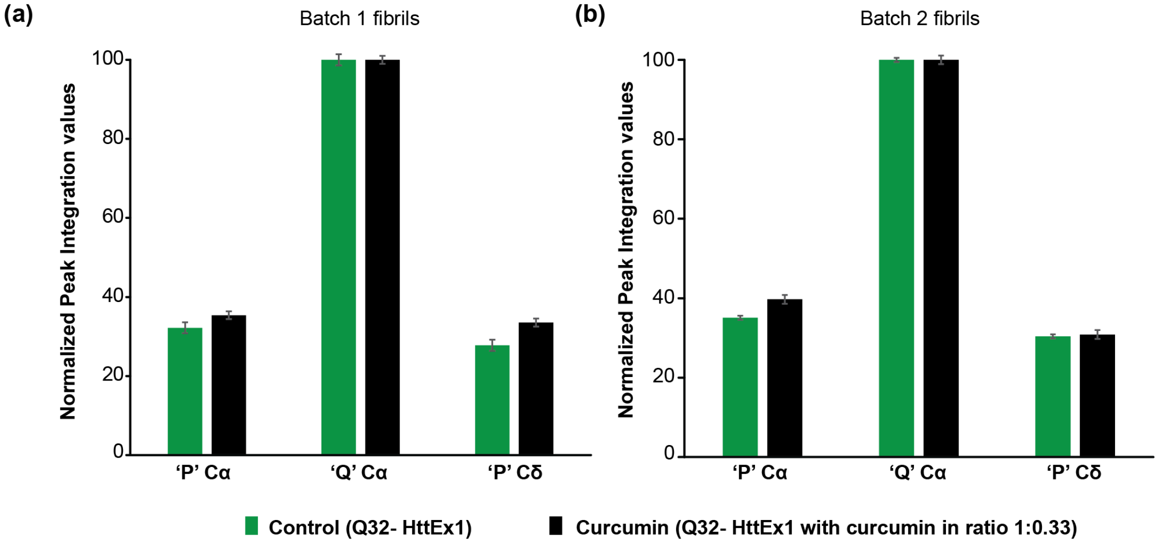


**Supplementary Fig. 8. SSNMR peak integration of Proline and Glutamine peaks of Q32-HttEx1 and Q32-HttEx1 with curcumin (1:0.33).** Bar graphs representing the normalized peak integration values of the proline and glutamine peaks shown in the $^{13}C$ CP spectra in (a) supplementary figure 6, batch 1 fibrils and (b) supplementary figure 7, batch 2 fibrils. Normalization of the peak areas was performed relative to the glutamine $C\alpha$ peaks (containing both the a and b type Gln conformers). Green bars correspond to values from the Q32-HttEx1 fibrils and black bars from the Q32-HttEx1 with curcumin (1:0.33). The error bars indicate an estimate of noise.

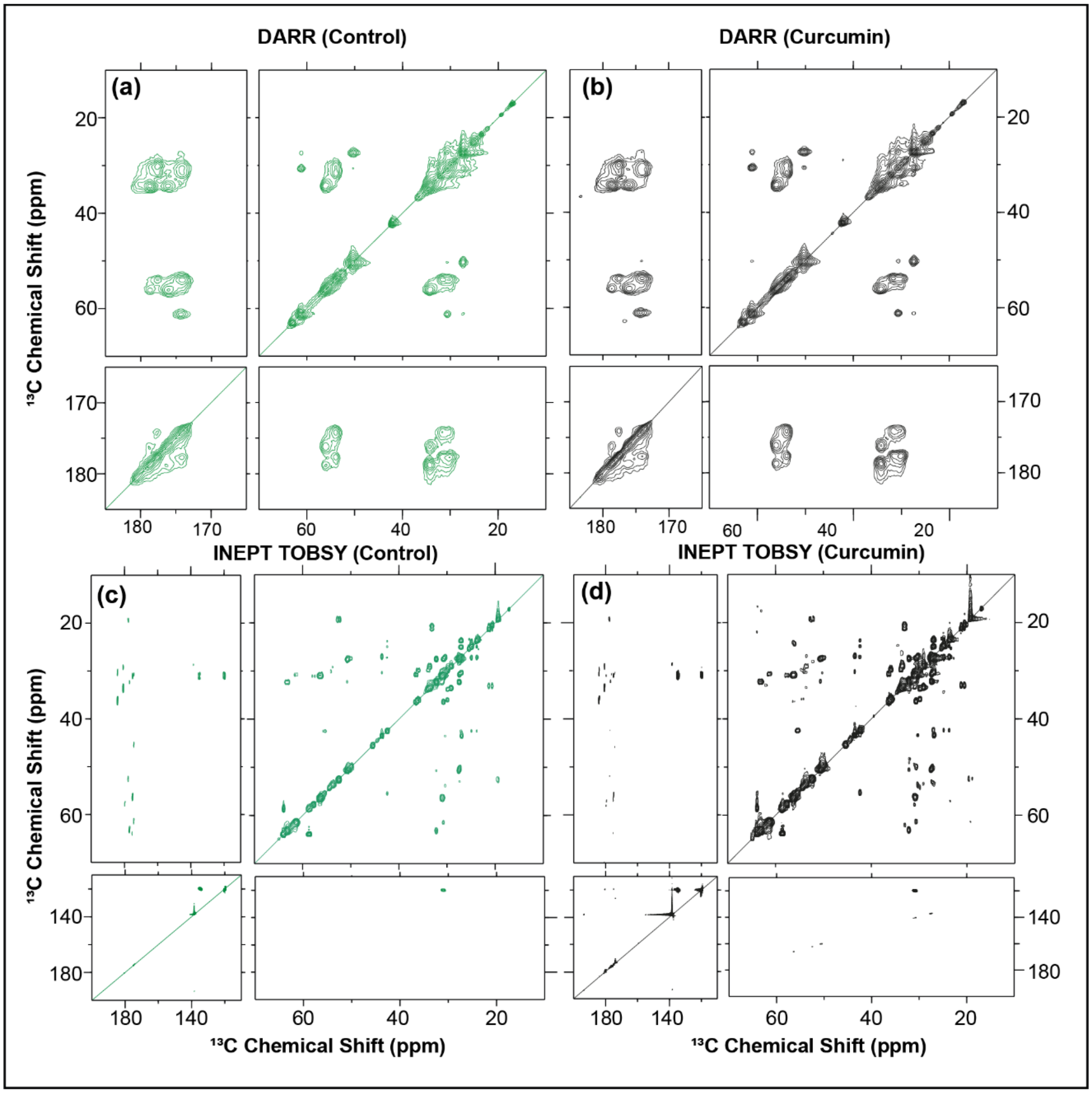


**Supplementary Fig. 9. 2D ssNMR analysis of Q32-HttEx1 fibrils formed in presence (black) and absence (green) of curcumin (Batch 2).** (a) $^{13}C$ -$^{13}C$ DARR spectrum for U-$^{13}C$, $^{15}N$ Q32-HttEx1 fibrils obtained at 13 kHz MAS and 25ms of DARR mixing. (b) $^{13}C$ -$^{13}C$ DARR spectrum for U-$^{13}C$, $^{15}N$ Q32-HttEx1 fibrils prepared in presence of curcumin (1:0.33) obtained at 13 kHz MAS and 25ms of DARR mixing. (c) $^{13}C$ -$^{13}C$ INEPT-TOBSY spectrum for U-$^{13}C$, $^{15}N$ Q32-HttEx1 fibrils obtained at 10 kHz MAS. (d) $^{13}C$ -$^{13}C$ INEPT-TOBSY spectrum for U-$^{13}C$, $^{15}N$ Q32-HttEx1 fibrils prepared in presence of curcumin (1:0.33) obtained at 10 kHz MAS.

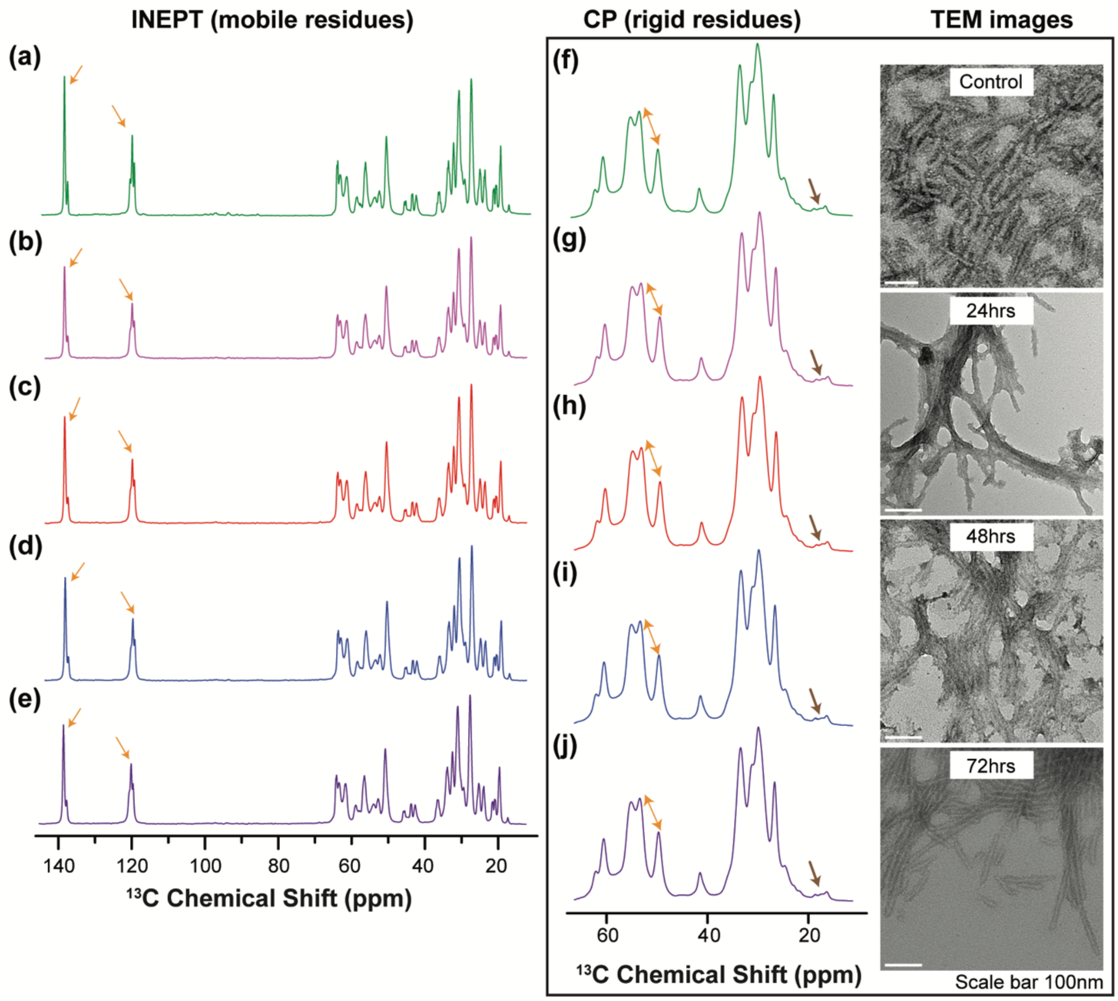


**Supplementary Fig. 10. MAS ssNMR and electron microscopy of curcumin addition after aggregation of Q32-HttEx1.** 1D INEPT $^{13}$C spectra of (a) Q32-HttEx1 fibrils without curcumin, (b) Q32-HttEx1 pre-formed fibrils treated with curcumin (1:0.33) for 24 hours, (c) 48 hours, (d) 72 hours, and (e) 96 hours. Zoomed 1D CP $^{13}$C spectra's of (f) Q32-HttEx1 fibrils without curcumin, (g) Q32-HttEx1 pre-formed fibrils treated with curcumin (1:0.33) 24 hours, (h) 48 hours, (i) 72 hours, and (j) 96 hours. TEM images of the pre-formed fibrils with and without treatment with curcumin (1:0.33) are shown on the right. Orange arrows in INEPT spectra indicate residues from the C-terminal tail showing small changes in intensity. The orange arrows in the CP panel mark the signal intensity differences between the glutamine and proline signals (compare to Figure 4). The brown arrows in the CP panel indicate peaks for Ala residues in Htt$^{NT}$ of the HttEx1 fibrils (2).

**Supplementary Table 1. Labeling scheme and amount of isotopically labeled MAS ssNMR samples.** In all cases, the indicated Q32-HttEx1 proteins were studied as mature amyloid- like fibrils that had been formed at room temperature. The reference to batches indicates two independently prepared samples (see Extended Methods).

| Name | Description | Labeling details | Sample size |
|---|---|---|---|
| Sample 1 | Q32-HttEx1 fibrils, Batch 1 | U- $^{13}C$, $^{15}N$ | 2.5mg |
| Sample 2 | Q32-HttEx1 inhibited with curcumin (1:0.33) fibrils, Batch 1 | U- $^{13}C$, $^{15}N$ | 2.5mg |
| Sample 3 | Q32-HttEx1 fibrils, Batch 2 | U- $^{13}C$, $^{15}N$ | 15mg |
| Sample 4 | Q32-HttEx1 inhibited with curcumin (1:0.33) fibrils, Batch 2 | U- $^{13}C$, $^{15}N$ | 20mg |
| Sample 5 | Q32-HttEx1 pre-formed fibrils treated with curcumin (1:0.33) | U- $^{13}C$, $^{15}N$ | 11.5mg |

**Supplementary Table 2. Detailed experimental conditions of the MAS NMR experiments**. Abbreviations: NS, number of scans per $t_1$ point; MAS, magic angle spinning rate; RD, recycle delay; TPPM, $^1$H decoupling power during evolution and acquisition (using two-pulse phase modulation scheme); $t_1$ evolution, number and length (in $\mu$s) of $t_1$ evolution increments. Temperature of the cooling gas for all the experiments was 275K and the mixing time for DARR experiments was 25ms. Sample identification: the sample details are mentioned in Supplementary Table 1 (above).

| **Figure*** | **Sample** | **Experiment** | **NS** | **MAS (kHz)** | **RD (s)** | **TPPM (kHz)** | **$t_1$ evol. ($\mu$s)** | **Contact time (ms)** |
|---|---|---|---|---|---|---|---|---|
| 4(a,b,e) S5 | Sample 1 | 1D $^{13}$C CP | 1024 | 10 | 3 | 83.3 | | 1 |
| S5 | Sample 1 | 1D $^{13}$C INEPT | 1024 | 10 | 3 | 50 | | |
| S5 | Sample 1 | 1D $^{13}$C HPDEC | 1024 | 10 | 3 | 83.3 | | 1 |
| 4(a,b,e) S5 | Sample 2 | 1D $^{13}$C CP | 1024 | 10 | 3 | 83.3 | | 1 |
| S5 | Sample 2 | 1D $^{13}$C INEPT | 1024 | 10 | 3 | 50 | | |
| S5 | Sample 2 | 1D $^{13}$C HPDEC | 1024 | 10 | 3 | 83.3 | | 1 |
| S6, S9 (f) | Sample 3 | 1D $^{13}$C CP | 1024 | 10 | 3 | 83.3 | | 1 |
| S6, S9 (a) | Sample 3 | 1D $^{13}$C INEPT | 1024 | 10 | 2.79 | 50 | | |
| S6 | Sample 3 | 1D $^{13}$C HPDEC | 1024 | 10 | 3 | 83.3 | | 1 |
| 4c S7(a) | Sample 3 | 2D DARR | 40 | 13 | 2.79 | 83.3 | 760x36 = 27360 | |
| S7(c) | Sample 3 | 2D INEPT-TOBSY | 32 | 8.33 | 3 | 70 | 640x30.11= 19270.4 | |
| S6 | Sample 4 | 1D $^{13}$C CP | 1024 | 10 | 3 | 83.3 | | 1 |
| S6 | Sample 4 | 1D $^{13}$C INEPT | 1024 | 10 | 2.79 | 50.6 | | |
| S6 | Sample 4 | 1D $^{13}$C HPDEC | 1024 | 10 | 3 | 83.3 | | 1 |
| 4(d) S8(b) | Sample 4 | 2D DARR | 40 | 13 | 2.79 | 83.3 | 760x36 = 27360 | |
| S8(c) | Sample 4 | 2D INEPT-TOBSY | 32 | 8.33 | 3 | 72 | 640x30.11= 19270.4 | |
| S9 (b-e) | Sample 5 | 1D $^{13}$C INEPT | 1024 | 10 | 3 | 50 | | |
| S9 (g-j) | Sample 5 | 1D $^{13}$C CP | 1024 | 10 | 3 | 83.3 | | 1 |

* *Figure identifiers with an S-number refer to figures in the Supplementary Information.*

**Supplementary Table 3. SAXS analysis parameters.** Fitted parameters using the equation discussed in methods section included in the Supplementary Information, above.

| | A | σ | $\bar{r}$(nm) | L* (nm) | d (nm) | $\upsilon$ | B | C | n |
|---|---|---|---|---|---|---|---|---|---|
| **Q32-HttEx1** | 6.7e-10 | 0.28 | 6.5 | 1000 | 24 | 100 | 2.9e-05 | 1.15e-03 | 2.26 |
| **Q32-HttEx1 with curcumin (1:0.33)** | 3.9e-09 | 0.35 | 4.5 | 1000 | 20 | 180 | 6.5e-04 | 1.4e-03 | 2.16 |

*this parameter was kept fixed to the mentioned value during the fitting

**Supplementary Table 4. Fitted parameters for the SAXS analysis using the model of Perevozchikova et al. (11)**. Parameters with subscript 2 refer to large objects/aggregates while 1 stands for small objects, the fibrils in our case.

| | $G_2$ | $R_{g2}$* | $S_2$* | $S_1$ | $R_{g1}$ (nm) | M |
|---|---|---|---|---|---|---|
| **Q32-HttEx1** | 0.05 | 1000 | 1 | 1 | 6 | 3 |
| **Q32-HttEx1 with curcumin (1:0.33)** | 0.045 | 1000 | 1 | 1 | 3.7 | 3 |

*these parameters were kept fixed to the mentioned value during the fitting
